# A novel method for monitoring ground-dwelling arthropods on hard substrates: characterizing arthropod biodiversity among survey methods

**DOI:** 10.1101/2021.12.06.471448

**Authors:** Katherine McNamara Manning, Kayla I. Perry, Christie A. Bahlai

## Abstract

Sampling approaches are commonly adapted to reflect the study objectives in biodiversity monitoring projects. This approach optimizes findings to be locally relevant but comes at the cost of generalizability of findings. Here, we detail a comparison study directly examining how researcher choice of arthropod trap and level of specimen identification affects observations made in small-scale arthropod biodiversity studies. Sampling efficiency of four traps: pitfall traps, yellow ramp traps, yellow sticky cards, and a novel jar ramp trap were compared with respect to an array of biodiversity metrics associated with the arthropods they captured at three levels of identification. We also outline how to construct, deploy, and collect jar ramp traps. Trapping efficiency and functional groups of arthropods (flying, crawling, and intermediate mobility) varied by trap type. Pitfalls and jar ramp traps performed similarly for most biodiversity metrics measured, suggesting that jar ramp traps provide a more comparable measurement of ground-dwelling arthropod communities to pitfall sampling than the yellow ramp traps. The jar ramp trap is a simple, inexpensive alternative when the physical aspects of an environment do not allow the use of pitfalls. This study illustrates the implications for biodiversity sampling of arthropods in environments with physical constraints on trapping, and the importance of directly comparing adapted methods to established sampling protocol. Future biodiversity monitoring schemes should conduct comparison experiments to provide important information on performance and potential limitations of sampling methodology.

## Introduction

There are many ways to observe populations and communities of insects. A vast literature of entomology studies aim to optimize trapping and monitoring methods for particular arthropod taxa and conservation goals (Agosti et al., 2000; Henderson & Southwood, 2016; Montgomery et al., 2021; O’Connor et al., 2019; Osborne et al., 2002). Specific trapping methods have been developed to reflect the arthropod community of interest as well as the physical or logistical constraints of the focal environment. Yet, this variability in sampling approach creates challenges for biodiversity monitoring. The effectiveness of conservation management programs is dependent on reproducible, reliable, and comparable data as these can impact biodiversity research outcomes, especially over time (Cardinale et al., 2018). Other challenges of biodiversity monitoring include errors in detection, misidentification, geographical constraints, and incomplete or biased views of the population or community (Saunders et al., 2019), especially as new survey formats are developed with technological advances and community science involvement (Isaac et al., 2020). Therefore, measurements of biodiversity are context-dependent, varying based on the methods used, and interacting with other elements from the environment that vary over time and space. Thus, the outcomes of biodiversity assessments that often inform conservation management strategies or policy are dependent on sampling methodology (Busse et al., 2022; Elphick, 2008; Gardiner et al., 2012; Prendergast et al., 2020; Saunders et al., 2019; Vallecillo et al., 2020; Whitworth et al., 2017).

Arthropod sampling methodology may be particularly prone to introducing contextual biases to data, which makes biodiversity monitoring difficult to approach in a comprehensive, standardized way (Montgomery et al., 2021). Each collection method has variable trapping efficiency that depends on arthropod biology and behavior as well as trap design (Montgomery et al., 2021). These biases do not eliminate the utility of the collected data, but additional information about the goals, constraints, and methods of a given experiment or monitoring strategy must be used to contextualize and understand the limitations and further use of these data. This contextual information also aids effective synthesis of data across biodiversity studies (Elphick, 2008). Within insect ecology, there is a strong cultural precedent of ‘do-it-yourself’ approaches for developing novel trapping methods, customized to a given situation (examples include: Bouchard et al., 2000; Dowd et al., 1992; Knuff et al., 2019; Owino, 2011; Russo et al., 2011; White et al., 2016). This customization tends to make the findings from arthropod surveys very adaptable, but results are also relatively contextually-specific. For example, insect traps are typically designed to catch a specific subset of a community, relevant to study goals. Sticky cards, flight intercept traps, and pan traps (also known as bee bowls) are all designed to catch flying insects. However, even among common sampling methods for flying insects, there is variation in trap design (for example, coloured pan/bowl trapping: Gonzalez et al., 2020; Joshi et al., 2015; Toler et al., 2005; Tuell & Isaacs, 2009; Vrdoljak & Samways, 2012).

Once samples are collected, further contextual biases may occur through the processing, identifying, and recording of arthropod biodiversity data. Because arthropods are numerically abundant and diverse, processing and identifying all specimens within samples can be a logistical challenge. The time and specialized taxonomic training required to identify arthropods beyond order or family level makes processing all samples to the species level an unrealistic goal for many studies. Depending on the study, researchers may address this challenge by focusing on a subset of individuals within a specific taxon or group of taxa. Alternatively, researchers may identify more individuals but at coarser taxonomic or functional classifications. This heterogeneity in the taxonomic resolution of arthropod data can make direct comparisons among studies difficult (Ferro & Summerlin, 2019) and has the potential to undermine ecological synthesis (Michener & Jones, 2012), but feasibility and goals of the study are still important to consider.

Use of common approaches may aid synthesis of monitoring data for arthropod populations, but may be constrained by the environments that these techniques are deployed in. For instance, pitfall traps are a commonly used method to sample ground-dwelling arthropods (Greenslade, 1964; Hohbein & Conway, 2018) and consist of a container filled with a killing fluid dug into the soil so that the rim is flush with the ground’s surface (Figure 1a). Although there are many benefits to using pitfall traps to sample ground-dwelling arthropods, there are several challenges and limitations. Importantly, there is not a standard trap design, material, or size for pitfall traps, which could impact syntheses across studies and global long-term monitoring of arthropod taxa (Brown & Matthews, 2016; Hohbein & Conway, 2018; Spence & Niemelä, 1994). Furthermore, some environments do not support the installation of pitfall traps to sample ground-dwelling arthropod communities, which may inhibit or bias biodiversity monitoring programs for these habitats. For example, thin-soil environments such as alvars, rocky glades, barrens, and green roofs have surface substrates that are too shallow to install conventional pitfall traps. Some biodiversity studies have employed an alternative ramp pitfall trap design which consists of a container placed on the ground with one to four ramps leading into the container (Bostanian et al., 1983; Bouchard et al., 2000, 2005; Patrick & Hansen, 2013; Weary et al., 2019). Abundant and diverse ground beetle communities were captured using ramp traps in alvar habitats in Ontario, Canada (Bouchard et al., 2005). Community composition of ground-dwelling beetles and spiders was similar among pitfall and ramp traps in oak woodland and chaparral habitats (Weary et al., 2019). However, similar to pitfall traps, ramp traps do not have a standard trap design, material, or size, and in some cases, may be challenging to build and transport due to trap size and complexity (Weary et al., 2019).

**Figure 1:**
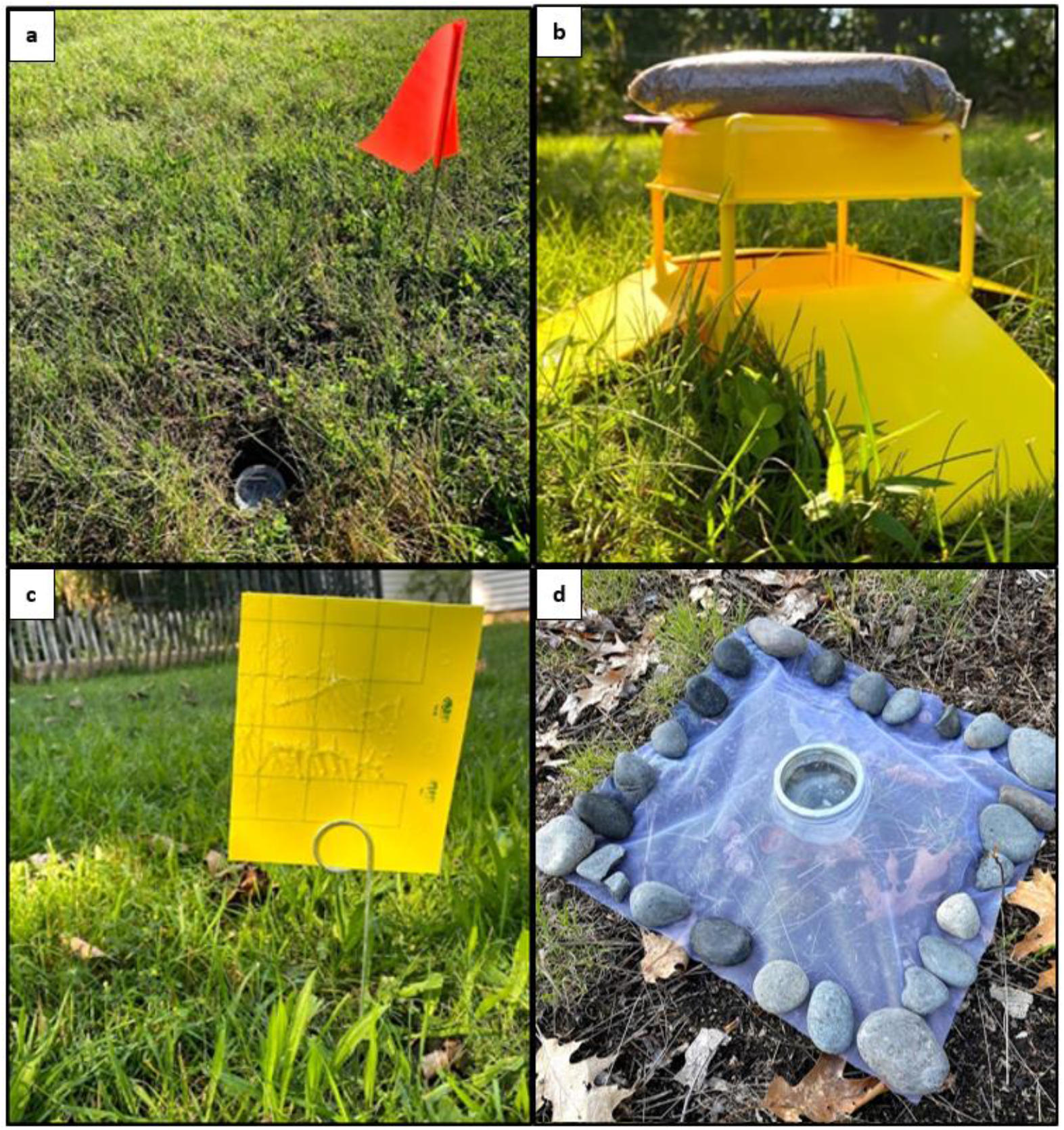
Arthropod traps: pitfall (a), yellow ramp trap (b), yellow sticky card (c), jar ramp trap (d) deployed at sampling sites in Kent, Ohio.

The objective of this study was to investigate how the design of arthropod traps affect the observations of arthropod communities, particularly for trap designs that had been adapted to contend with physical constraints of their deployment environment. Specifically, we compared the performance of two traps designed to minimize disruption to soil substrates to two classical trapping methods. We compared arthropod communities among traditional pitfall traps, commercially available ramp traps, sticky cards, and a novel, alternative design to the commercial ramp trap, the jar ramp trap. Herein, we outline how to construct and deploy jar ramp traps. We predicted that the arthropod community captured by each trap would vary based on the structure of the trap and the functional biology of the arthropods. In addition to comparing these four sampling methods, we compared how multiple approaches to insect identification may impact the findings. We predicted that different identification levels will produce variable statistical results, each revealing and obscuring different parts of the community, and suggesting tradeoffs between both trapping and sample processing approaches. We discuss recommendations for comparison studies which will improve the interoperability of data produced by specialized insect sampling methodology.

## Materials and Methods

### Study sites

We selected study sites with similar abiotic attributes to the thin-soil environments our adaptive traps were designed for: exposed to solar radiation, precipitation, and wind, but with deeper soils to accommodate the use of pitfall traps. We selected three mown horticultural grasslands in Northeast Ohio in the City of Kent, owned by Kent State University and operated by the Kent State University Center for Ecology and Natural Resource Sustainability. No pesticide, herbicide, or fertilizer was directly applied to any of the sampling locations for at least 12 months prior to our study.

### Arthropod sampling

At each site, four trap types were deployed to sample arthropod communities: 1) pitfall traps; 2) commercially available yellow ramp traps; 3) commercially available yellow sticky cards; and 4) novel jar ramp traps. Two replicates of each trap type were installed 3-5 m apart at each site for 48 hours during a period of warm, dry weather every other week during the months of July, August, and September 2020, amounting to seven sampling periods and 21 location-date replicates in total. All traps used in this study have a common bias in that they only detect active arthropods that move into the trap rather than extract individuals from a given area of habitat. As with all passive trapping methods, samples are not a measurement of raw abundance, per se, but instead a measure of activity density. These captures are a good proxy for population abundance if activity rates are density independent (Didham et al., 2020).

Pitfall traps consisted of a 100 ml transparent plastic specimen container, 7.5 cm in height with a 4.5 cm diameter opening, filled with soapy water (Dawn Original Liquid Dish Soap, Procter & Gamble, Cincinnati, OH, USA) (using similar methodology to Cates et al., 2021; Sultaire et al., 2021; Wills et al., 2019) (Figure 1a). Yellow ramp traps (ChemTica Internacional S.A., Santo Domingo, Costa Rica) were square yellow plastic containers (14 × 14 × 13 cm) with a roof and detachable ramps (30% slope) on four sides, placed on the ground’s surface and filled with soapy water (Figure 1b). A small sandbag (sand inside quart zipper-top bag) was placed on top of the roof to minimize movement of the trap in windy conditions. These ramp traps are commercially available but have not been extensively tested in the field to sample ground-dwelling arthropod communities. In a survey of North American Great Lakes Basin thin-soil environments our group observed high numbers of flying insects in these commercially available ramp traps, while characteristic ground-dwelling arthropod taxa were absent. Yellow sticky cards (Pherocon, Zoecon, Palo Alto, CA, USA) were cut in half to limit disturbance by wind (11×14 cm) and affixed to wire stands, positioning the top of the card approximately 30 cm off the ground (Figure 1c). Because the commercial ramp trap collected primarily flying insects in our previous survey, we included sticky cards in the trap comparison to examine any overlap in community composition with the ground-dwelling arthropod traps. Jar ramp traps were constructed using a 41 cm x 41 cm square of noseeum mesh attached to the rim of an open, shallow clear glass Ball jar (Ball Corporation, Broomfield, CO, USA) (236 ml, 7.5 cm diameter, 5 cm height) filled with soapy water, with small rocks or stones to secure the mesh to the ground (Figure 1d). The jar ramp trap was engineered to address some structural issues with the commercial trap, improve the sample handling experience, and collect arthropod communities that more closely match a pitfall trap.

Trap contents were collected after 48 hours. Samples from the yellow ramp traps were strained with noseeum mesh in the field and preserved in 70% ethanol in gallon plastic zipper-top bags. Yellow sticky cards were placed directly into gallon plastic zipper-top bags. Jar ramp traps and pitfall traps had plastic lids secured on the glass jar or plastic container, respectively, directly in the field. Samples were processed in the laboratory and specimens were identified with the aid of a dissecting microscope. For the duration of the study, yellow sticky cards were stored in the freezer and all other samples in vials with 70% ethanol.

In contrast to the commercial yellow ramp trap, the ramps on the jar ramp traps were at a lower angle and the noseeum mesh provided a substrate that was easier to grip than the smooth plastic. Additionally, the design of the jar ramp trap improved handling and sample collection in the field, as a plastic lid easily seals the sample jar in the field until sampling processing in the laboratory. Construction of the jar ramp trap began by cutting a square of mesh (approximately 41 by 41 cm). The lid was removed from the Ball jar, flat piece discarded, and the screw band reattached. A thin layer of glue (Gorilla Heavy Duty Construction Adhesive, Gorilla Glue, Cincinnati, OH, USA) was applied around the top of the screw band (Figure 2b). The mesh square was placed over the opening of the jar and screw band, with the jar in the center of the mesh, and secured to the screw band by applying light pressure (Figure 2c). This was left to dry for the recommended time on the glue instructions. Then, using a utility knife, we cut out the mesh on the inside of the jar opening. This left us with a plain glass jar and a detachable “ramp” (Figure 2d). To deploy the trap, we attached the mesh ramp to the Ball jar, adjusting so that the edges were flat. The outer edges of the mesh were lined with small stones. The Ball jar was filled to the top with soapy water: a gallon of water with about 1 teaspoon of Dawn Original dish soap (Figure 1d). For trap collection, the stones were removed, then the mesh ramp was gently unscrewed from the jar and a plastic lid was screwed onto the jar. The samples could now be transported, with light padding, and temporarily stored for up to one week (Figure 2e). Jars, ramps, and plastic lids were easily washed and reused.

**Figure 2:**
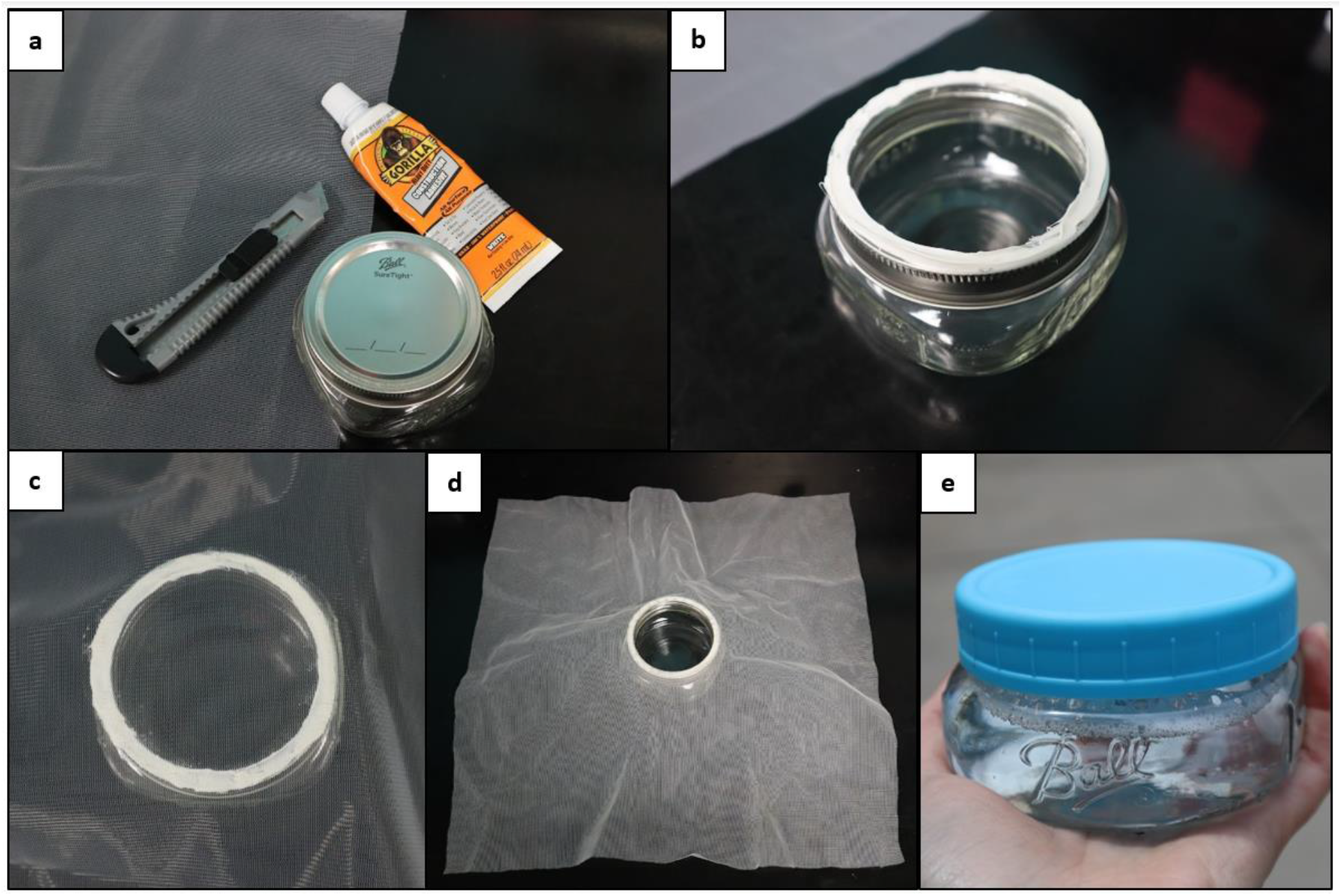
Step by step photos of jar ramp trap construction.

### Arthropod identification

To examine the impact of specimen identification approach on observations, specimens were identified using three approaches: taxonomic order, functional classification, and focal taxa to species/genus. To conduct the order-level classification, all arthropod specimens collected were determined to their taxonomic order using Borror & White, 1998 and Marshall, 2017. For functional classification, arthropods were categorized based on primary mobility: flying, crawling, or intermediate (Borror & White, 1998; Evans, 2014; Marshall, 2017). Intermediate means that they are ground-dwelling arthropods that may have the capability of flying (i.e. Carabidae, Staphylinidae), have wings but primarily jump (i.e. Cicadellidae, Orthoptera), or adults that may or may not have wings (i.e. Aphididae). Specimens were identified to order (Acarina, Araneae, Collembola, Diptera, Lepidoptera, Orthoptera, Thysanoptera, Zygoptera), superfamily (Apoidea, Chalcidoidea, Ichneumonoidea), group (“wingless parasitiod wasps”), or family. This was modeled after studies that used this mixed approach of identifying for other insect functional classifications such as natural enemy or herbivore (Fiedler & Landis, 2007; Gibson et al., 2019), or predators (Hermann et al., 2019; Mabin et al., 2020). For the focal taxa approach, specimens captured from three functionally-important beetle (Order Coleoptera) families were identified to the highest taxonomic resolution possible (genus or species). For these determinations, we focused on individuals from Carabidae (Lindroth, 1961-1969), Staphylinidae (Brunke et al., 2011; Klimaszewski et al., 2018), and Coccinellidae (Evans, 2014; Gardiner et al., 2006; Gardiner, 2015).

### Statistical analyses

All statistical analyses were completed using R 4.1.3 (R Core Team, 2022). Unless otherwise noted, all analyses were performed at each level of arthropod identification (taxonomic order, functional classification, and focal taxa to species/genus). Data were evaluated for statistical assumptions of normality and homogeneity of variance. Accumulation curves for each trap type were created using the *BiodiversityR* package (Kindt & Coe, 2005). To estimate sampling efficiency for each trap type, we used nonparametric Jackknife order 1 estimator to compare observed and estimated richness. Abundance (number of arthropods per trap), taxonomic richness (number of taxa per trap), Shannon diversity index (Hill, 1973), and Pielou’s evenness index (Pielou, 1966) were calculated using the *vegan 2*.*5-7* package (Oksanen et al., 2019).

Generalized linear mixed effects models (GLMM) were developed using the *lme4* (Bates et al., 2015) and *lmerTest* (Kuznetsova et al., 2017) packages to examine differences in arthropods among trap types. The response variables examined were arthropod abundance, richness, diversity, and evenness. Each GLMM included trap type and sampling date as categorical fixed effects and trap number nested within the site as a random effect. The global model took the form: *Response variable ∼ Trap + Date + (1*|*Site:Replicate)*. For response variables involving count data, the Poisson family error structure was initially specified. Models examining arthropod abundance at the order and functional level failed to meet model assumptions and were given the negative binomial error structure instead. The categorical fixed effect date was excluded from GLMMs examining functional level arthropod diversity and evenness to meet model assumptions. The function ‘Anova’ from the *car* package (Fox & Weisberg, 2019) was used to examine significance of trap type in each model. Tukey pairwise comparisons were performed using the *emmeans 1*.*7*.*4-1* package (Lenth, 2021) for all models.

For the functional classification level of identification, we also performed a functional group analysis in which specimens were classified into groups of crawling, flying, or intermediate mobility (excluding groups where insufficient identification prevented assigning a functional role) to assess differences in abundance and richness by trap type using generalized linear models with the form: *Response variable ∼ Trap*. Negative binomial error structure was used for abundance models because they failed to meet normality assumptions. Similar to the GLMMs, models were developed using the *lme4* and *lmerTest* packages, and Tukey pairwise comparisons were performed using the *emmeans 1*.*7*.*4-1* package.

To characterize the arthropod communities collected by each of the four trap types, we used non-metric multidimensional scaling (NMDS, with Bray-Curtis distance). Permutational multivariate analysis of variance (PERMANOVA), analysis of multivariate homogeneity of group dispersions (BETADISPER), and pairwise multilevel comparison using Adonis were performed following each NMDS analysis to assess compositional dissimilarity between trap types. NMDS, PERMANOVA, and BETADISPER were computed using functions in the *vegan 2*.*5-7* package. Pairwise adonis was performed using the *pairwiseAdonis* package (Martinez Arbizu, 2020).

## Results

Seven sampling periods at our three sites yielded 165 samples (accounting for three pitfalls lost to disturbance by mammal excavation), which contained a total of 13,634 arthropod specimens. Overall, yellow ramp traps caught the greatest number of individuals (7,758); followed by sticky cards (4,199); then jar ramp traps (1,099); with pitfall traps catching the least (578) (see abundances by identification level: Table S1). Trap types had openings of various sizes (Figure S1). The capture of functional groups of arthropods (flying, crawling, or intermediate) and individual taxonomic groups varied by trap type.

### Order-level analysis

The total number of orders captured varied from 9 (pitfall traps) to 12 (yellow ramp traps and jar ramp traps) (Figure 3a). When compared with first order jackknife richness estimates, pitfall trap efficiency was 90%; yellow ramp trap efficiency was 100%; yellow sticky card efficiency was 100%; and jar ramp trap efficiency was 86%.

**Figure 3:**
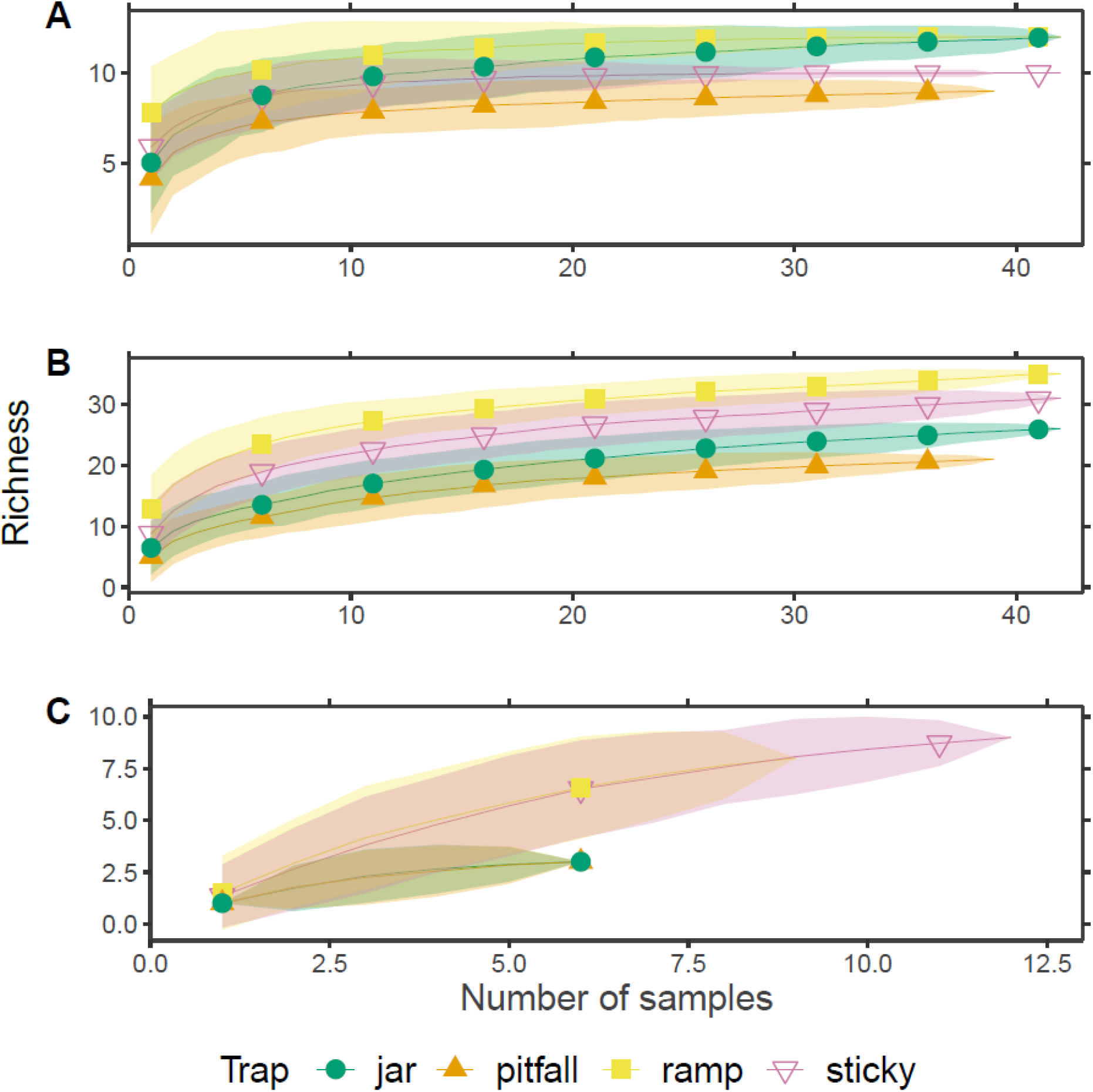
Taxon richness accumulation curves for each trap type by identification level: (A) order, (B) functional, (C) focal taxa.

Overall differences in richness, abundance, Shannon diversity, and evenness were observed between the trap types in generalized linear mixed effect models (Table S2; Figure 4a). Higher arthropod richness, abundance, and diversity were observed in yellow ramp traps than other trap types. For richness and abundance, yellow ramp traps were followed by yellow sticky traps, jar ramp traps, and pitfall traps, all of which differed statistically from each other. Compared to yellow ramp traps, all other trap types captured similar levels of arthropod diversity. Pitfall traps and jar ramp traps had high arthropod evenness compared to yellow ramp traps and sticky cards.

**Figure 4:**
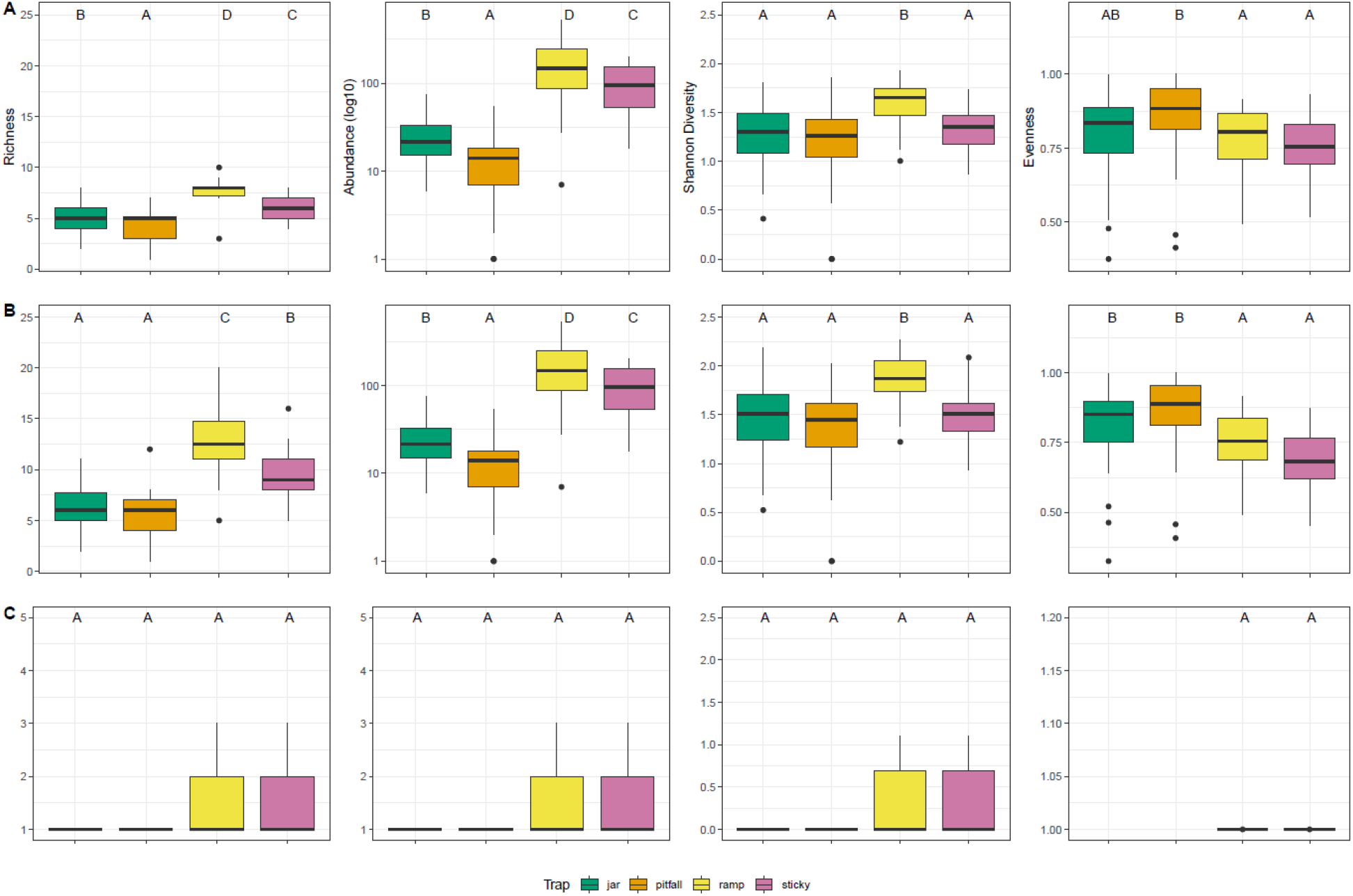
Richness, abundance (log 10), Shannon diversity, and evenness between trap types on the identification level of: (A) Order, (B) Functional, (C) Focal taxa. Letters shared indicate no statistical difference in estimated marginal means by Tukey method, P <0.05.

Community composition varied between all trap types at the order level (p = 0.001, Figure 5a). Homogeneity of multivariate dispersion could not be assumed, indicating that some trap types had more variable community composition than others.

**Figure 5:**
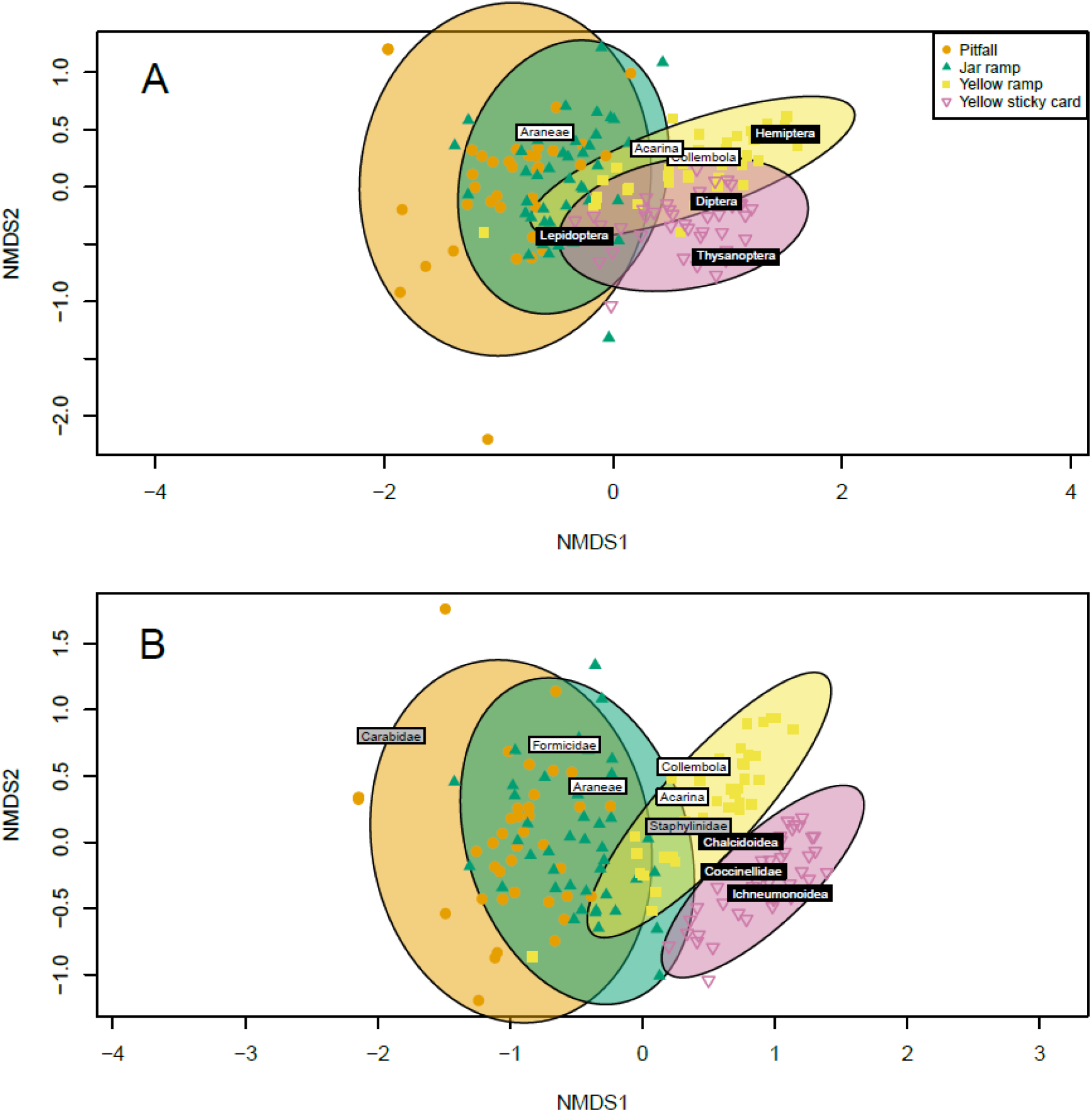
Non-metric multidimensional scaling (NMDS) of arthropod community composition in each of four trap types deployed in managed grasslands in Kent, Ohio (USA) in 2020. (A) Order: stress = 0.14, (B) functional: stress = 0.15. Ellipsoids represent 95% confidence of the mean for trap types. Points displayed represent community composition for each trap type. Select flying guild taxa displayed as white text on black boxes, background and ground-crawling guild taxa displayed as black text on in white boxes, background and intermediate in grey boxes.

### Functional-level analysis

The total unique taxa captured varied from 21 (pitfalls) to 35 (yellow ramp traps) (Figure 3b). The flying group consisted of 22 taxa with a mean abundance across all trap types of 43.2 ± 16.3 and mean richness of 3.9 ± 0.9. The crawling group consisted of five taxa with a mean abundance across all trap types of 18.4 ± 8.1 and mean richness of 2.6 ± 0.5. The intermediate group consisted of 10 taxa with a mean abundance across all trap types of 20.5 ± 15.9 and mean richness of 1.7 ± 0.3 (Table S1). When compared with first order jackknife richness estimates, pitfall trap efficiency was 84%; yellow ramp trap efficiency was 84%; yellow sticky card efficiency was 84%; and jar ramp trap efficiency was 79%.

Overall differences in richness, abundance, Shannon diversity, and evenness were observed between the trap types in generalized linear mixed effect models (Table S2; Figure 4b). Similar to the order level analyses, yellow ramp traps collected the highest richness, abundance, and diversity. Jar ramp traps and pitfall traps collected the lowest richness. For abundance, yellow ramp traps were followed by yellow sticky cards, jar ramp traps, and pitfall traps, all of which differed statistically from each other. Compared to yellow ramp traps, all other trap types captured similar levels of diversity. Pitfall traps and jar ramp traps had the highest evenness, followed by yellow ramp traps and sticky traps.

Community composition varied between all trap types when taxa were grouped by functional classification (p = 0.001, Figure 5b). As with the order-level analysis, homogeneity of multivariate dispersion could not be assumed.

Arthropod abundance and richness in each trap type were then compared by functional group (flying, crawling, or intermediate) (Table S3; Figure 6). Sticky cards and yellow ramp traps captured the highest abundance and richness of flying arthropods. Yellow ramp traps captured the highest abundance and richness of crawling arthropods. Yellow sticky cards captured the lowest abundance and richness of crawling arthropods. Yellow ramp traps captured the highest abundance and richness of intermediate mobility arthropods. Richness of intermediate mobility arthropods was low overall, and captures were fairly consistent among other trap types.

**Figure 6:**
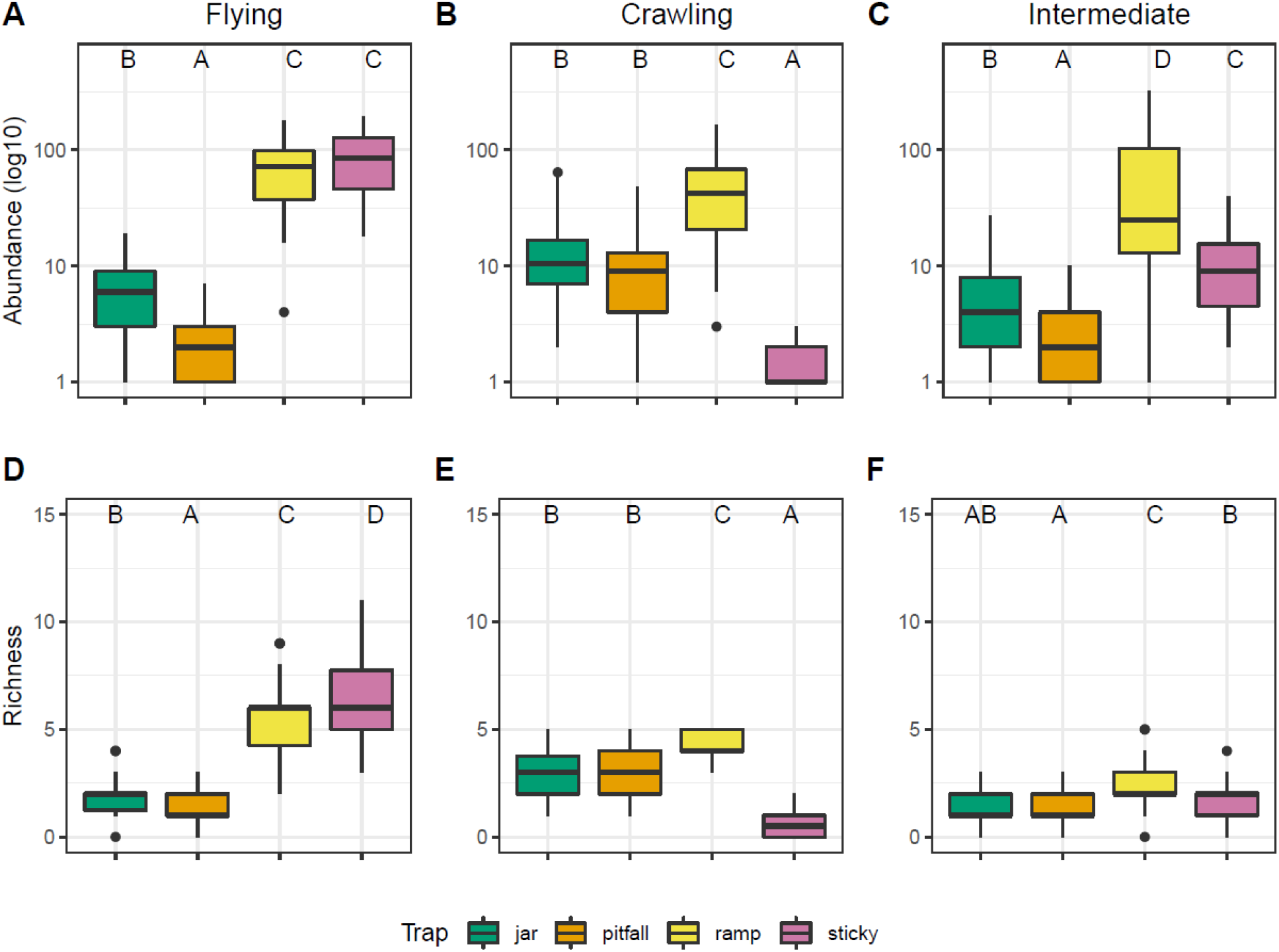
Flying, crawling, and intermediate mobility arthropod abundance and richness by trap type. (A) flying abundance, (B) crawling abundance, (C) intermediate abundance, (D) flying richness, (E) crawling richness, (F) intermediate richness. Note that abundance was log10 transformed. Letters shared indicate no statistical difference in estimated marginal means by Tukey method, P <0.05.

### Focal taxon analysis

Six specimens comprising three species of Carabidae were collected: *Cicindelidia punctulata* (Olivier), *Cratacanthus dubius* (Palisot de Beauvois), and *Harpalus faunus* (Say). Twenty-one adult specimens (larvae were not identified to species) comprising seven species of Coccinellidae were collected: *Brachiacantha ursina* (Fabricius), *Coleomegilla maculata* (Degeer), *Cycloneda munda* (Say), *Harmonia axyridis* (Pallas), *Hippodamia variegata* (Goeze), *Hyperapis undulata* (Say), and *Propylea quatuordecimpunctata* (Linnaeus). Staphylinidae were identified to genus. Sixteen specimens comprising five genera were identified: *Acrotona, Meronera, Rabigus, Stenus*, and *Xantholinus*. Four staphylind specimens were damaged and could not be identified.

The total number of focal taxa captured varied from 3 (pitfalls and jar ramp traps) to 9 (yellow sticky cards) (Figure 3c). When compared with first order jackknife richness estimates, pitfall trap efficiency was 78%; yellow ramp trap efficiency was 69%; yellow sticky card efficiency was 77%; and jar ramp trap efficiency was 78%.

There were no differences in richness, abundance, Shannon diversity, or evenness detected between trap types in generalized linear mixed effects models (Table S2; Figure 4c), likely due to sparse data.

Community composition varied by trap type (p = 0.003), but only some trap comparisons were statistically significantly different: pitfall traps and yellow sticky cards and jar ramp traps and yellow sticky cards. However, because data was sparse, NMDS results failed to converge. Homogeneity of multivariate dispersion was assumed.

## Discussion

Ultimately, biodiversity trends observed in any study are highly sensitive to sampling methodology (Berglund & Milberg, 2019; Brice et al., 2021; Gardiner et al., 2012; Joshi et al., 2015; O’Connor et al., 2019; Prendergast et al., 2020; Whitworth et al., 2017). For example, in a European bumble bee survey the three methods used all produced different estimates of the population (Wood et al., 2015). The results of the present study are no exception. Even between methods specifically adapted to particular habitat structures, traps captured different arthropod communities. However, our study also suggests an important caveat: the ability to detect differences between sampling types is also affected by sample size. Studies that focus at a high taxonomic resolution will require many more individual samples to be able to detect differences. Our analyses of focal taxa were not able to detect statistical trends due to the relatively small number of specimens from each group.

In our study, the jar ramp trap and pitfall trap communities were very similar, suggesting they had similar performance when deployed in the environment. For both the order and functional classification levels, no differences were observed among jar ramp traps and pitfall traps for the majority of the biodiversity metrics. When broken down by functional mobility groups, jar ramp and pitfall traps had similar richness and abundance of crawling arthropods, which is their target mobility group. Pitfall traps are commonly and widely used to sample ground-dwelling arthropods (Greenslade, 1964; Southwood, 1978), and these findings suggest that jar ramp traps are a suitable alternative design. These two trap types also had similar intermediate mobility richness, a functional group which includes ground-dwelling arthropods that walk or run on the soil surface, but may be capable of flight, such as beetles in the families Carabidae and Staphylinidae (Larochelle & Larivière, 2003; Levesque & Levesque, 1995).

For example, in a study of ground beetles in thin-soiled alvar environments, 91% of species captured had fully developed hind wings, making them capable of flight (Bouchard et al., 2005). Analyses of community composition among jar ramp and pitfall traps indicated large overlap at the order and functional level, further supporting the comparable measurement of these trap types. Jar ramp traps should be considered when pitfall sampling cannot be used, for example in areas with shallow soils that do not allow for pitfall traps to be placed into the ground.

Though ramp traps are an alternative method to sample the ground-dwelling arthropod community without disturbing the substrate (Bouchard et al., 2000; Patrick & Hansen, 2013; Weary et al., 2019), we found that yellow ramp traps were not a sufficient alternative to pitfall traps. In a preliminary study using the yellow ramp traps as our primary means of collecting ground-dwelling arthropods, we observed one carabid individual, and had initially concluded that these bioindicators (Koivula, 2011; Rainio & Niemela, 2003; Serap & Luff, 2010) were rare at our sample sites. In this study, yellow ramp traps captured a community of arthropods dominated by flying taxa that was more similar to yellow sticky cards than pitfall traps or jar ramp traps. Yellow sticky cards, or glue traps, are a commonly used methodology for sampling flying populations of insects, especially in agricultural ecosystems (Aliakbarpour & Rawi, 2011; Bahlai et al., 2015; Gardiner et al., 2009; Li et al., 2021; Muppudathi et al., 2018; Musters et al., 2021). The bright yellow color of the yellow ramp trap may result in a similar attractiveness to flying arthropods as observed for yellow sticky cards (Shimoda & Honda, 2013; Shin et al., 2020). The yellow ramp trap captured a higher abundance, richness, and Shannon diversity than the jar ramp trap, pitfall trap, or yellow stick card in the order and functional level analyses. This pattern is likely due to the yellow ramp trap collecting ground-dwelling and flying arthropods, which is supported by the overlap in community composition among these trap types. Therefore, these numbers may be misleading, as the trap was designed to sample ground-dwelling arthropods, and thus may not be an authentic measurement of function in ground-dwelling communities when it is catching the flying community as well. Traps capturing a higher number of non-target arthropods may obscure biodiversity trends associated with a study’s goals depending on the level of taxonomic resolution. For instance, in a native bee survey in an agroecosystem in Pennsylvania, blue vane traps captured the greatest richness and abundance, however, they were trapping higher ratios of common bees to rare bees compared to the pan traps used in the study (Joshi et al., 2015).

Our findings highlight the contextual dependence of insect sampling methodologies. Although the commercial yellow ramp traps were designed to collect ground-dwelling arthropods, these traps have often been used in conjunction with a chemical lure. For example, in the literature these yellow ramp traps are most commonly used to target large, ground-dwelling pest weevil species in agricultural landscapes (Oehlschlager et al., 2002; Reddy et al., 2008; Reddy et al., 2009). Because yellow ramp traps are used with a lure in these circumstances, the biology and behavior of target pest taxa were able to overcome the structural issues that these traps presented to other arthropods when used in the context of passive trapping. Although yellow ramp traps are commercially available and do capture a variety of ground-dwelling and flying arthropods, considerations should be made about the goals of the study before employing these traps to passively sample arthropod communities.

Having an alternative trap for ground-dwelling insects is a necessity in situations where researchers are physically constrained from using pitfall traps. The novel jar ramp trap is inexpensive, easy and quick to construct, and simple to deploy, even in comparison to other homemade ramp traps (Bouchard et al., 2000; Patrick & Hansen, 2013; Weary et al., 2019). The plastic lids make sample collection and transport very user-friendly. The yellow ramp trap required users to remove the four ramps from the trap, drain the contents, and transfer them to another container with ethanol in the field. This was cumbersome and created opportunities for specimen loss. Additionally, removal and reattachment of ramps often broke the connection point on the trap. On the jar ramp trap, noseeum mesh served as a 360 degree ramp around the collection jar. With the aid of the rocks, the mesh ramp blends in relatively well in the environment. The mesh ramp creates a coarse surface which insects appear to have no trouble crawling on, compared to slick plastic ramps of the commercial traps, and the breathable, light colored material may create less of a change in microclimate than the yellow plastic on hot days.

Though the jar ramp trap has many advantages over the yellow ramp trap, and performed similarly to pitfall traps, it does have limitations. The Ball jar is only 5 cm deep, so leaving the traps deployed for an extended period of time may result in issues such as evaporation of the collection fluid or flooding in the case of heavy rainfall. Larger jars could easily be adapted to this design, however, at the compromise of ramp steepness. Although the rocks provide a natural means of securing the noseeum mesh to the ground, the presence of rocks could affect movement of some ground-dwelling arthropods. In spaces where some substrate exists, the traps may be lightly covered by soil to secure the mesh.

Our study demonstrated that the level to which taxa are identified impacts the study results and researcher interpretation of the biodiversity data. In this study, arthropods were identified using three approaches (i.e. taxonomic order, functional classification of mobility, and focal taxa to genus/species), and each level of identification provided different information about the arthropod communities. For example, clear patterns among trap types were observed at the order and functional levels of identification, but there was insufficient data at the focal taxa level. The order level may be the most comparable across biodiversity monitoring studies because it requires less taxonomic expertise and fewer samples to reach high sampling efficiency. However, this coarse level of identification can miss information that may be vital to the goals of the study. In this study, identification to order did not allow us to examine whether the trap types were capturing ground-dwelling arthropods because orders contain species of diverse functional mobility groups. For example, Order Hymenoptera is composed of wasps and bees, which fly, and ants, most of which crawl. In contrast, a relatively small number of specimens were collected of the focal beetle taxa, which resulted in the lower estimated sampling efficiency of 79%. Although sampling efficiency at the species level was lower than at the order level, this estimate is comparable to other studies that investigate arthropod communities using species level taxonomic resolution. Ground beetles collected using unbaited pitfall traps in greenspaces within nearby Cleveland, Ohio documented 69% of the estimated species richness in one year of study and 66% the next (Perry et al., 2020).

Classification of arthropods by functional mobility groups provided an additional dimension of biodiversity data that harnessed ecological life history information for each taxonomic group. The functional classification level of identification provided a compromise between the relative speed at which samples could be enumerated, similar to the order level classification, but with more meaningful interpretation of results based on arthropod biology. This classification scheme allowed us to examine captures among trap types based on their primary mobility, which facilitated our understanding of how trap design influences the observations of arthropod communities. Importantly, the use of functional traits to study patterns of biodiversity has several advantages. Functional traits provide a stronger connection to ecosystem processes and function than taxonomic measures of diversity such as abundance and diversity (Gagic et al., 2015), and have a greater comparative applicability across habitats and ecosystems (Webb et al., 2010). Therefore, functional classifications provide a complementary approach to traditional taxonomic metrics that can improve biodiversity monitoring programs with minimal additional effort.

Although the focal beetle taxa required larger sample sizes to detect differences that may exist among trap types, we still observed some members of the important predatory beetle families Carabidae, Staphylinidae, and Coccinellidae as well as which trap types captured them (Table S1). This allowed us to dive deeper into two ground-dwelling, intermediate mobility families (Carabidae and Staphylinidae) and one flying family (Coccinellidae). Three species within the family Carabidae were collected during this study. *C. punctulata* and *H. faunus* are macropterous (i.e. have fully developed wings), and thus, associated with greater dispersal ability as this trait renders them capable of flight (Larochelle & Larivière, 2003). *C. dubius* is wing-dimorphic, meaning individuals can be either macropterous or brachypterous (i.e. reduced wings, incapable of flight) (Larochelle & Larivière, 2003).

Interestingly, each ground-dwelling trapping method caught a different species: pitfall traps caught all three *H. faunus*; a yellow ramp trap caught the single *C. dubius*; and jar ramp traps caught the two *C. punctulata*. Five genera of Staphylinidae were identified in the study. The predatory genus *Stenus* has large bulbous eyes, is uniquely diurnal, and can move on water by the release of an alkaloid from their abdomen (Evans, 2014; Gardiner, 2015).

Compared to species of Carabidae, staphylinid genera were more evenly spread among the different trap types, but yellow ramp traps and yellow sticky cards captured the majority of individuals. Only one specimen each of *Rabigus* (in yellow ramp), *Stenus* (in yellow ramp), and *Xantholinus* (on yellow sticky card) were observed. The remaining and most abundant individuals belonged to *Acrotona* (in yellow and jar ramp traps) and *Meronera* (all trap types). Seven species of Coccinellidae were captured during the study, with over 70% of the lady beetle specimens collected by yellow sticky cards, which are considered reliable traps for measuring activity density in this family (Bahlai et al., 2013). Yellow ramp traps captured some lady beetles as well, but neither the pitfall nor the jar ramp trap captured this taxon reliably.

It is not uncommon for biodiversity monitoring to occur in sensitive habitats with unique constraints, requiring customized approaches to monitoring. However, these modifications to standardized trapping methods limit the comparability of study findings. This study illustrates the implications for biodiversity sampling of arthropods in environments with physical constraints on trapping, and the importance of directly comparing adapted methods to established sampling protocol. We have shown that conducting a comparison of those methods can provide important contextual information on how that method performs, and its potential limitations in monitoring protocol.

Comparison studies should ideally be conducted in the environment where monitoring will occur. However, in our case the thin-soil environments that jar ramp traps and yellow ramp traps are meant for did not allow for the use of pitfall traps. By conducting the comparison in an environment with similar abiotic attributes as thin-soil sites, we were able to comprehensively examine the efficacy of these trap types to inform future arthropod monitoring study designs in these sensitive habitats. This study leverages sites that were accessible and relatively uniform in environmental conditions to demonstrate that such comparisons of methodology can be relatively small scale and accomplished with limited labor. Indeed, the experimental work for this study was completed on a university campus when travel and support labor was highly limited by the COVID-19 lockdown.

Despite its importance to environmental management, developing standards for biodiversity monitoring comes with many challenges. Between idiosyncratic biology of target taxa and habitat effects on trapping efficiency, and indeed, trap structure, it becomes essential to compare modified trapping methodology against standards to ensure transferability of data. Future biodiversity monitoring schemes, especially those occurring in sensitive or unusual habitats, should conduct comparison experiments to maximize the chances of capturing target taxa, while minimizing disturbance to their habitat and thus, activity patterns of taxa, as well as fostering future ecological synthesis.

## Acknowledgments

Data used in our study was collected on the ancestral and contemporary lands of the Erie and Seneca peoples where Kent, Ohio is currently located. The Kent State University Center for Ecology and Natural Resource Sustainability and their director Lauren Kinsman-Costello allowed for the use of properties to conduct this research. Kent State University Facilities Management, especially the Care of Grounds Crew and their director Rebekkah Berryhill coordinated grounds care around our sampling. Particular thanks to Julia Perrone, MLIS for assistance in trap and experimental design and Timothy D. Niepokny for assistance in the field. This work was completed with the support of a grant from the National Science Foundation Faculty Early Career Development Program from the Division of Biological Infrastructure (DBI 2045721) to CB.

## Author Contributions

KMM and CB conceived research. KMM conducted experiments. KMM and KP analyzed data and conducted statistical analyses with the aid of CB. KMM wrote the article with critical revisions by CB and KP. CB secured funding. All authors read and approved the manuscript.

## Data Availability

Data and relevant code for this research work are stored in GitHub: https://github.com/katiemmanning/jar_ramp_trap and have been archived within the Zenodo repository: https://doi.org/10.5281/zenodo.6994127.

---

**Figure S1:**
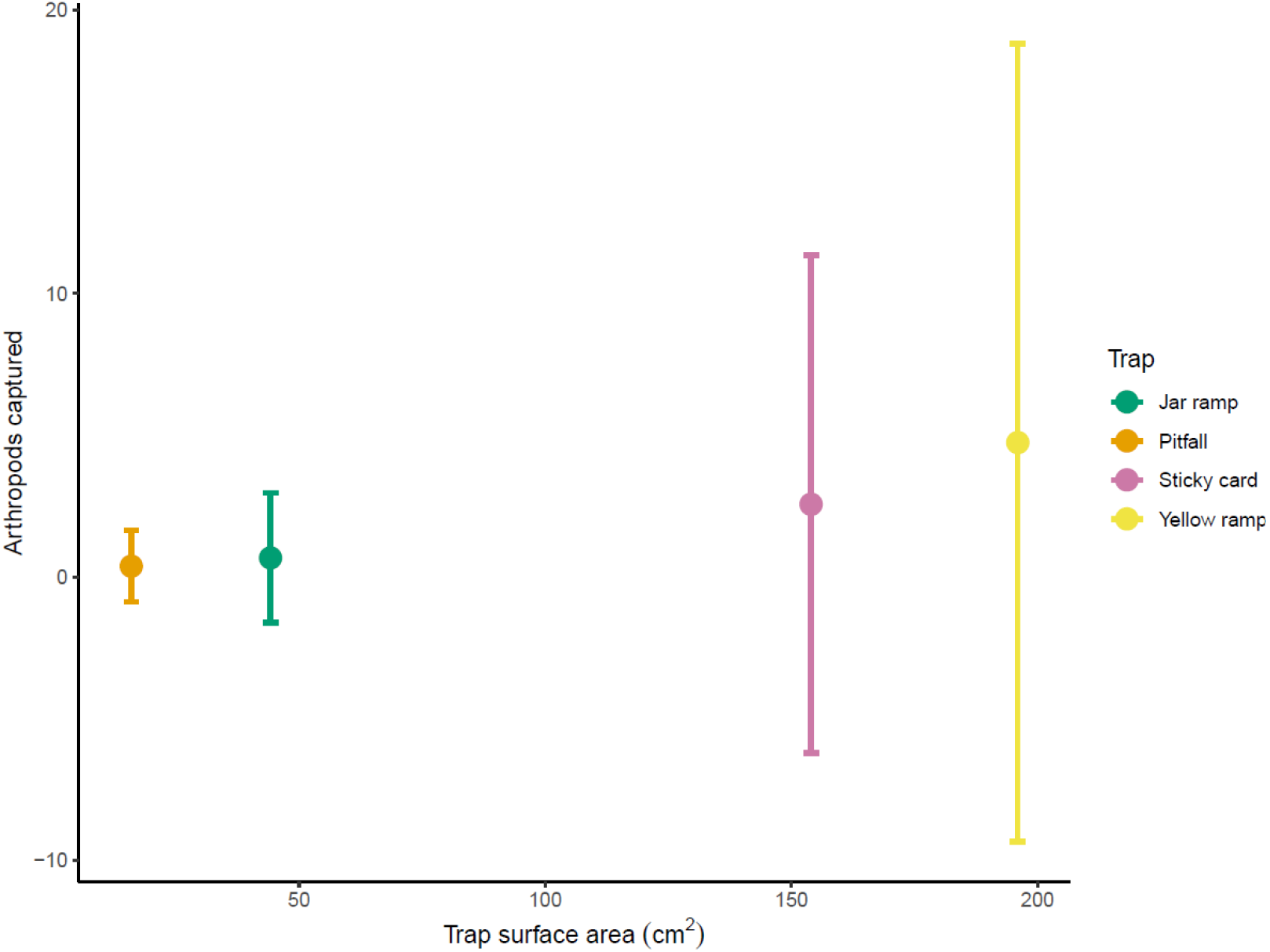
Arthropods captured in each trap type by surface area of the trap opening. The variability of arthropods captured and mean catch increased with trap surface area.

